# Variable inhibition of unwinding rates of DNA catalyzed by the SARS-Cov-2 (COV19) helicase nsp13 by structurally distinct single DNA lesions

**DOI:** 10.1101/2021.10.13.464299

**Authors:** A.H. Sales, S. Ciervo, T. Lupoli, V. Shafirovich, N.E. Geacintov

## Abstract

The SARS 2 (Covid 19) helicase nsp13 plays a critically important role in the replication of the Corona virus by unwinding double-stranded RNA (and DNA) with a 5’⟶3’ strand polarity. Here we explored the impact of single, structurally defined covalent DNA lesions on the helicase activity of nsp13 in aqueous solutions, The objectives were to derive mechanistic insights into the relationships between the structures of DNA lesions, the DNA distortions that they engender, and the inhibition of helicase activity. The lesions included two bulky stereoisomeric N2-guanine adducts derived from the reactions of benzo[a]pyrene diol epoxide with DNA. The *trans*-adduct assumes a minor groove conformation, while the *cis*-product adopts a base-displaced intercalated conformation. The non-bulky DNA lesions included the intra-strand cross-linked thymine dimers, the *cis-syn*-cyclobutane pyrimidine dimer, and the pyrimidine (6−4) pyrimidone photoproduct. All four lesions strongly inhibit the helicase activity of nsp13, The UV photolesions feature a 2 - 5-fold smaller inhibition of the nsp13 unwinding activity than the bulky DNA adducts, and the kinetics of these two pairs of DNA lesions are also different. The connections between the structural features of these four DNA lesions and their impact on nsp13 unwinding efficiencies are discussed.

## Introduction

The lifecycle of the SARS-CoV-2 virus and its replication in infected human host cells depends critically on its mechanism of replication. The nsp13 helicase belongs to the NSF1 superfamily of helicases that plays a critical role in the replication of the virus by first unwinding double-stranded RNA to provide a single-stranded template for the transcription of the viral genome. The reduction of functional helicase activities by small molecule inhibitors has been considered for potential cancer therapy applications.^*1, 2*^ More recently, the COVID19 pandemic stimulated significant interest in the design of new inhibitors^*3–5*^ for suppressing the SARS-Cov-2 helicase unwinding activities and the replication of the virus.^*6*^

The nsp13 helicase unwinds unmodified double-stranded RNA^*6, 7*^ or DNA^*6–10*^ with a 5’⟶ 3’ translocation polarity by an ATP-driven mechanism. The helicase can be inhibited either by blocking ATP hydrolysis,^*7*^ or by inhibiting the unwinding of the nsp13 helicase without necessarily affecting the ATPase activity.^*10–15*^ The latter approach has received more attention than the direct inhibition of the unwinding mechanism.^*4, 16–18*^

The impact of non-covalent binding of various small molecules on the unwinding of double-stranded DNA by various helicases has been studied.^*19*^ However, connections between helicase unwinding efficiencies and the molecular structures of the non-covalent DNA complexes are difficult to establish; furthermore, the potential formation of structurally different complexes cannot be excluded. Thus, the impact of single, covalent, and structurally defined DNA lesions on unwinding activities could provide useful insights into helicase umwinding structure-function relationships.^*19, 20*^ However, this approach has received much less attention, and thus remains an understudied area in this field.

In this communication, we summarize preliminary results that indicate that the UV cross-linked thymine dimer photolesions, the cyclobutane pyrimidine thymine dimer (CPD) and the pyrimidine (6−4) pyrimidone photoproduct, 6-4)PP (Fig. 1), and the bulky polycyclic aromatic DNA adducts (Fig. 2), diminish unwinding of double-stranded DNA with variable efficiencies. The two bulky DNA adducts (Fig. 1) were derived from the *cis- or trans*-addition *of* the exocyclic amino group of guanine to the C10 atom of the aromatic diol epoxide derivative (+)-7*R*,8*S*)-dihydroxy-(9*S*,10*R*)-epoxy-7,8,9,10 tetra-hydrobenzo[*a*]pyrene ((+)-*anti*-B[*a*]PDE).^*21, 22*^ The UV photolesions, the cyclobutane pyrimidine dimer (CPD) and the pyrimidine(6−4)pyrimidone photoproducts, were generated by UV light, and separated and purified by HPLC methods.^*23*^ The structural features of the two stereoisomeric B[*a*]PDE-*N*^2^-dG adducts positioned in double-stranded DNA duplexes have been established by high-resolution NMR methods (Fig. 2). The *trans* adduct is positioned externally in the minor groove of DNA oriented towards the 5’-direction of the modified strand.^*22*^ The stereoisomeric *cis* adduct is intercalated, and the modified guanine residue is displaced into the minor groove, while its partner cytosine is displaced into the major groove^*21*^ (Fig. 2). Analysis of high resolution NMR data, indicates that Watson-Crick base pairing is absent at the (6-4)PP crosslinked TT step and at the adjacent base pair on the 3’-side, and that the DNA duplex is bent by a rigid 44^0^ bend.^*24*^ By contrast, hydrogen bonding at the TT CPD dimer is maintained with a much smaller bend in the duplex of only ~9^0^.^*25*^

**Fig. 1.**
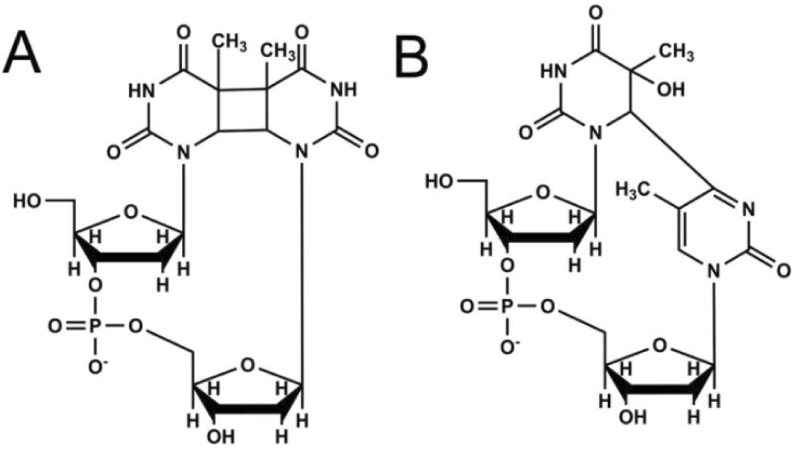
(A) Structures of CPD, and (B) (6 - 4)PP.

**Fig. 2.**
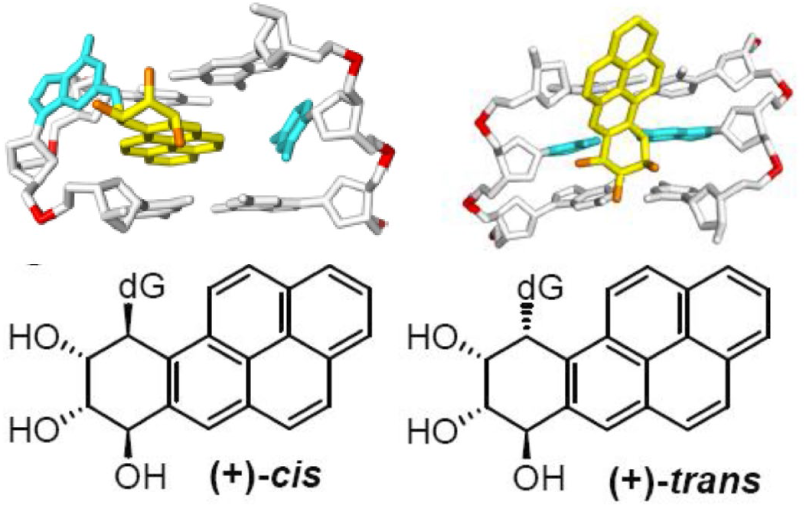
Stereochemical features and conformations of B[*a*]PDE-*N*^2-^dG adducts.

## Methods

The oligonucleotide substrate used in the unwinding assays is shown below:

> 5’-GCTTGCATGCCTGCAGGTCGACTCTAGAGGATCCATC**X**CTACCTACGAGG-BHQ2 3’-TCTCCTAGGTAGCGATGGATGCTCC-Cy3

The 3’-ends of the top, 5’---> 3’ helicase translocation strands, were labeled with the BHQ2 fluorescence quencher, while the 5’-end of the bottom strand contained the fluorescent Cy3 dye opposite BHQ2. The top strand contained 25 base pairs in the double-stranded region, and 25 nucleotides in the single-stranded overhang. The DNA lesions are denoted by **X**. In the double-stranded DNA substrates, the fluorescence emission of Cy3 (λ_max_=564 nm) was generated by excitation with a green diode laser (515 nm). In the case of the fully double-stranded DNA sequence, the fluorescence of Cy3 is fully quenched by BHQ2 in the opposite strand. The unwinding of the double-stranded DNA substrates induced by the nsp13 helicase dissociates the two strands, and the fluorescence of Cy3 is not quenched. The kinetics of unwinding were thus determined by monitoring the Cy3 fluorescence intensity at 564 nm.^*26*^

At the 5 nM DNA concentration employed in these experiments, the re-association rate of the two separated strands was negligible on the time scale of the experiment. Prior to the unwinding experiments, the DNA substrates were pre-equilibrated with the nsp13 protein (60 nM), and the unwinding reactions were initiated by mixing this solution with aliquots of the unwinding buffer that contained ATP. All experiments were conducted at 25 ^0^C.

### Unwinding phenomena

Typical unwinding results are depicted in Fig. 2. All four DNA lesions were subjected to identical experimental conditions (DNA and protein concentrations, etc.) In general, the unwinding kinetics follow the classic exponential kinetics:

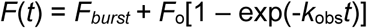

This equation features a burst of unwinding (*F*_*burst*_) that is associated with the rapid unwinding induced by pre-existing non-covalent helicase-DNA duplexes. The slower phase is due to subsequent complex formation and unwinding kinetics. Burst kinetics are indeed observed in the case of unmodified DNA (Fig. 3), but the burst is absent in the case of DNA containing any of the four DNA lesions (Fig. 4). The DNA lesions inhibit the burst activity, while the rates of unwinding depend on the nature of the DNA lesions and are ~ 20-100 times slower than in the case of unmodified DNA (Fig. 5). It is well established that the formation of oligomeric nsp13 complexes is necessary for successful unwinding activity,^*8*^ but it is not yet clear whether the DNA lesions inhibit the formation of productive oligomeric nsp13 complexes, or whether such complexes are formed but are unproductive in the presence of DNA lesions.

**Fig. 3.**
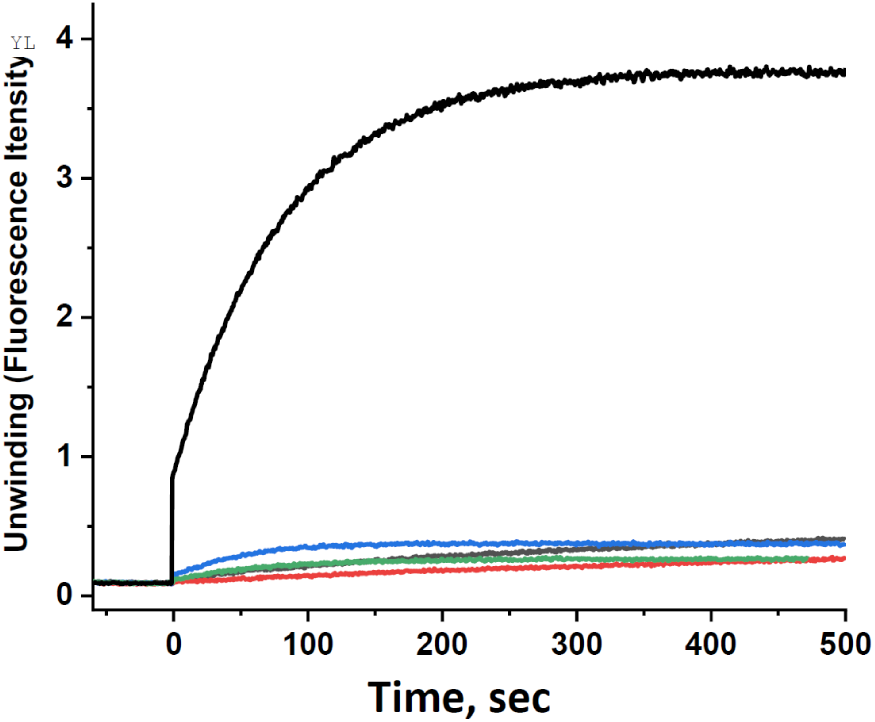
Unwinding of unmodified DNA by nsp13 (top). DNA lesions are shown on the bottom (same color scheme as in Fig. 4)

**Fig. 4.**
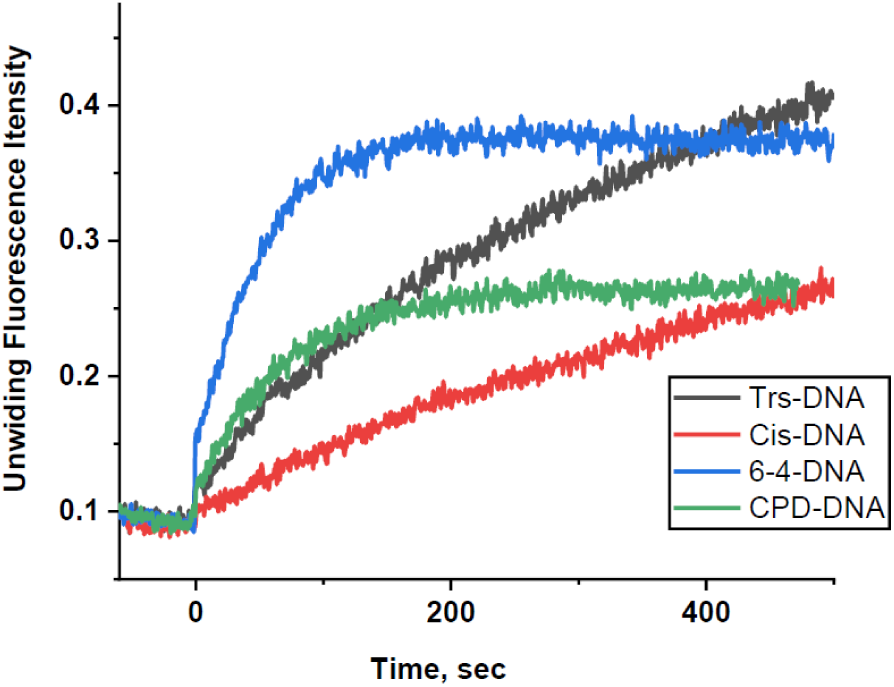
Impact of DNA lesions on the unwinding of double-stranded DNA by the nsp13 helicase.

**Fig. 5.**
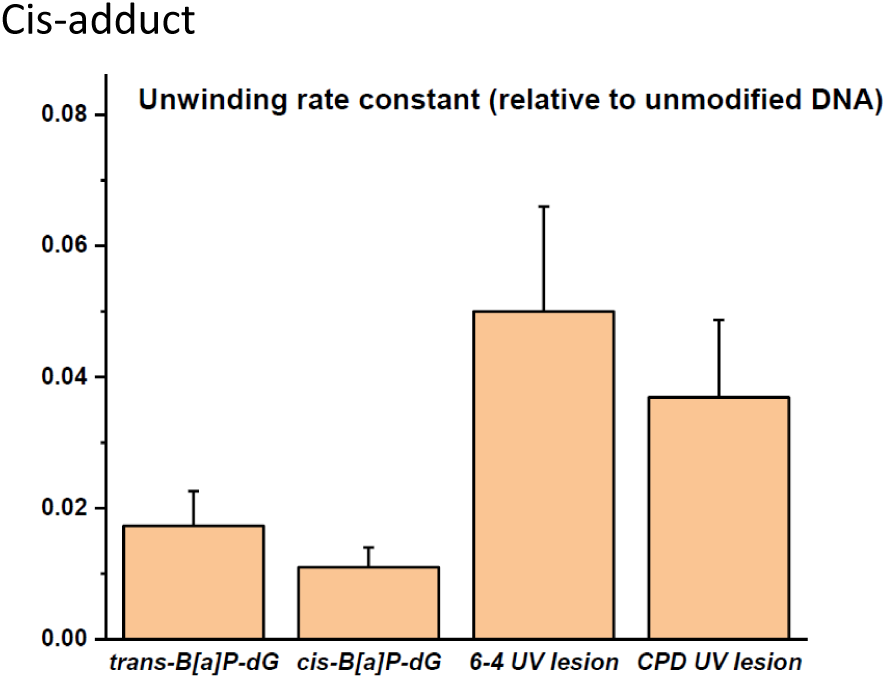
Unwinding rates relative to unmodified DNA. Averages of 5-7 independent experiments.

The differences in unwinding rates between the *cis-* and *trans-*BPDE-dG adducts suggest that the minor groove *trans* adduct^*22*^ is a lesser obstacle to unwinding than the intercalated base-displaced *cis* adduct.^*21*^ The *trans* adduct is characterized by intact base pairing at the site of the modified guanine and an almost normal backbone, while in the case of the *cis-*adduct the backbone is more distorted around the lesion site and the modified guanine is completely displaced from the interior of the duplex.^*22*^ In the case of the *trans-*adduct, the modified guanine residue remains hydrogen bonded to its partner strand residue C in the complementary strand, and the bulky aromatic ring system is located in the minor groove pointing towards the 5’-end of the modified strand.^*22*^ For successful unwinding to occur, the translocating strand must pass through a narrow channel formed by three of the five major domains of nsp13.^*27*^ However, the bypass of the bulky DNA lesions is only 2 - 5 times slower than the rates of unwinding of the ‘non-bulky thymine dimers CPD and (6-4)PP.

These results suggest that the intrastrand cross-linked DNA lesions CPD and (6-4)PP, although ‘non-bulky’, also strongly inhibit the unwinding of DNA catalyzed by nsp13. The CPD thymine dimer features two covalent cross-links rather than only one in (6-4)PP, but is overall less distorting than the (6-4)PP lesion.^*24, 25*^ The latter features a strong kink at the site of the lesion and a greater loss of base pairing than CPD, all of which is consistent with its moderately slower unwinding rate relative to CPD (Fig. 4).

Finally, the unwinding kinetics of the UV photolesions and the bulky B[*a*]PDE-*N*^2^-dG adducts are qualitatively different (Fig. 4). In the case of the UV lesions, the yields of unwinding products level off after reaching yields of 25 – 35%, while the rate of formation of products continues linearly up to yields of at least 25-35% in the case of the B[*a*]PDE-*N*^2^-dG adducts. Ongoing and future studies are focused on detailed analyses of the effects of nsp13 concentrations and oligomerization on binding equilibria and unwinding kinetics. These studies will include structurally different, bulky DNA lesions, and oxidatively derived guanine lesions^*28*^ that feature only one damaged nucleotide in the translocated strand rather than the two adjacent, intrastrand crosslinked thymine bases in CPD and (6-4)PP.

## Acknowledgements

We thank Hong Mu and Suse Broyde for supplying the DNA structures in Fig. 1. We also thank the Kapoor Lab at Rockefeller University for supplying the pET28(a)+ Nsp13 plasmid.^*27*^

## References

[1] Datta, A., and Brosh, R. M., Jr. (2018) New Insights Into DNA Helicases as Druggable Targets for Cancer Therapy, Front Mol Biosci 5, 59.

[2] Newman, J. A., and Gileadi, O. (2020) RecQ helicases in DNA repair and cancer targets, Essays Biochem 64, 819–830.

[3] Shyr, Z. A., Gorshkov, K., Chen, C. Z., and Zheng, W. (2020) Drug Discovery Strategies for SARS-CoV-2, J Pharmacol Exp Ther 375, 127–138.

[4] Xiu, S., Dick, A., Ju, H., Mirzaie, S., Abdi, F., Cocklin, S., Zhan, P., and Liu, X. (2020) Inhibitors of SARS-CoV-2 Entry: Current and Future Opportunities, J Med Chem 63, 12256–12274.

[5] Mengist, H. M., Dilnessa, T., and Jin, T. (2021) Structural Basis of Potential Inhibitors Targeting SARS-CoV-2 Main Protease, Front Chem 9, 622898.

[6] Romano, M., Ruggiero, A., Squeglia, F., Maga, G., and Berisio, R. (2020) A Structural View of SARS-CoV-2 RNA Replication Machinery: RNA Synthesis, Proofreading and Final Capping, Cells 9.

[7] Jang, K. J., Jeong, S., Kang, D. Y., Sp, N., Yang, Y. M., and Kim, D. E. (2020) A high ATP concentration enhances the cooperative translocation of the SARS coronavirus helicase nsP13 in the unwinding of duplex RNA, Sci Rep 10, 4481.

[8] Lee, N. R., Kwon, H. M., Park, K., Oh, S., Jeong, Y. J., and Kim, D. E. (2010) Cooperative translocation enhances the unwinding of duplex DNA by SARS coronavirus helicase nsP13, Nucleic Acids Res 38, 7626–7636.

[9] Ivanov, K. A., Thiel, V., Dobbe, J. C., van der Meer, Y., Snijder, E. J., and Ziebuhr, J. (2004) Multiple enzymatic activities associated with severe acute respiratory syndrome coronavirus helicase, J Virol 78, 5619–5632.

[10] Adedeji, A. O., and Lazarus, H. (2016) Biochemical Characterization of Middle East Respiratory Syndrome Coronavirus Helicase, mSphere 1.

[11] Shadrick, W. R., Ndjomou, J., Kolli, R., Mukherjee, S., Hanson, A. M., and Frick, D. N. (2013) Discovering new medicines targeting helicases: challenges and recent progress, J Biomol Screen 18, 761–781.

[12] Keum, Y. S., and Jeong, Y. J. (2012) Development of chemical inhibitors of the SARS coronavirus: viral helicase as a potential target, Biochem Pharmacol 84, 1351–1358.

[13] Yu, M. S., Lee, J., Lee, J. M., Kim, Y., Chin, Y. W., Jee, J. G., Keum, Y. S., and Jeong, Y. J. (2012) Identification of myricetin and scutellarein as novel chemical inhibitors of the SARS coronavirus helicase, nsP13, Bioorg Med Chem Lett 22, 4049–4054.

[14] Lee, C., Lee, J. M., Lee, N. R., Jin, B. S., Jang, K. J., Kim, D. E., Jeong, Y. J., and Chong, Y. (2009) Aryl diketoacids (ADK) selectively inhibit duplex DNA-unwinding activity of SARS coronavirus NTPase/helicase, Bioorg Med Chem Lett 19, 1636–1638.

[15] Cho, J. B., Lee, J. M., Ahn, H. C., and Jeong, Y. J. (2015) Identification of a Novel Small Molecule Inhibitor Against SARS Coronavirus Helicase, J Microbiol Biotechnol 25, 2007–2010.

[16] Gurung, A. B. (2020) In silico structure modelling of SARS-CoV-2 Nsp13 helicase and Nsp14 and repurposing of FDA approved antiviral drugs as dual inhibitors, Gene Rep 21, 100860.

[17] Ahmad, S., Waheed, Y., Ismail, S., Bhatti, S., Abbasi, S. W., and Muhammad, K. (2021) Structure-Based Virtual Screening Identifies Multiple Stable Binding Sites at the RecA Domains of SARS-CoV-2 Helicase Enzyme, Molecules 26.

[18] Shen, L., Niu, J., Wang, C., Huang, B., Wang, W., Zhu, N., Deng, Y., Wang, H., Ye, F., Cen, S., and Tan, W. (2019) High-Throughput Screening and Identification of Potent Broad-Spectrum Inhibitors of Coronaviruses, J Virol 93.

[19] Brosh, R. M., Jr., and Matson, S. W. (2020) History of DNA Helicases, Genes (Basel) 11.

[20] Khan, I., Sommers, J. A., and Brosh, R. M., Jr. (2015) Close encounters for the first time: Helicase interactions with DNA damage, DNA Repair (Amst) 33, 43–59.

[21] Cosman, M., de los Santos, C., Fiala, R., Hingerty, B. E., Ibanez, V., Luna, E., Harvey, R., Geacintov, N. E., Broyde, S., and Patel, D. J. (1993) Solution conformation of the (+)-cis-anti-[BP]dG adduct in a DNA duplex: intercalation of the covalently attached benzo[a]pyrenyl ring into the helix and displacement of the modified deoxyguanosine, Biochemistry 32, 4145–4155.

[22] Cosman, M., de los Santos, C., Fiala, R., Hingerty, B. E., Singh, S. B., Ibanez, V., Margulis, L. A., Live, D., Geacintov, N. E., Broyde, S., and et al. (1992) Solution conformation of the major adduct between the carcinogen (+)-anti-benzo[a]pyrene diol epoxide and DNA, Proc Natl Acad Sci U S A 89, 1914–1918.

[23] Kropachev, K., Kolbanovskii, M., Cai, Y., Rodriguez, F., Kolbanovskii, A., Liu, Y., Zhang, L., Amin, S., Patel, D., Broyde, S., and Geacintov, N. E. (2009) The sequence dependence of human nucleotide excision repair efficiencies of benzo[a]pyrene-derived DNA lesions: insights into the structural factors that favor dual incisions, J Mol Biol 386, 1193–1203.

[24] Kim, J. K., and Choi, B. S. (1995) The solution structure of DNA duplex-decamer containing the (6-4) photoproduct of thymidylyl(3’-->5’)thymidine by NMR and relaxation matrix refinement, Eur J Biochem 228, 849–854.

[25] Kim, J. K., Patel, D., and Choi, B. S. (1995) Contrasting structural impacts induced by cis-syn cyclobutane dimer and (6-4) adduct in DNA duplex decamers: implication in mutagenesis and repair activity, Photochem Photobiol 62, 44–50.

[26] Lohman, T. M., Tomko, E. J., and Wu, C. G. (2008) Non-hexameric DNA helicases and translocases: mechanisms and regulation, Nat Rev Mol Cell Biol 9, 391–401.

[27] Jia, Z., Yan, L., Ren, Z., Wu, L., Wang, J., Guo, J., Zheng, L., Ming, Z., Zhang, L., Lou, Z., and Rao, Z. (2019) Delicate structural coordination of the Severe Acute Respiratory Syndrome coronavirus Nsp13 upon ATP hydrolysis, Nucleic Acids Res 47, 6538–6550.

[28] Shafirovich, V., Kropachev, K., Anderson, T., Liu, Z., Kolbanovskiy, M., Martin, B. D., Sugden, K., Shim, Y., Chen, X., Min, J. H., and Geacintov, N. E. (2016) Base and nucleotide excision repair of oxidatively generated guanine lesions in DNA, J. Biol. Chem. 291, 5309–5319.

